# The Breast Cancer Epigenomics Track Hub

**DOI:** 10.1101/2022.05.01.490187

**Authors:** Giovanna Ambrosini, Andrea Agnoletto, Cathrin Brisken, Philipp Bucher

## Abstract

**Background:** Pioneering research has shown that high-throughput epigenomics assays such as ChIP-seq and ATAC-seq are applicable to patient-derived breast tumor samples. A host of public data has been accumulated since then, which are potentially of high value for basic research as well as personalized medicine. Such data sets constitute encyclopedias of biological knowledge. However, their impact has so far been limited by access obstacles, especially with regard to extraction and visualization of small portions of data that could potentially answer specific questions arising in a research context.

**Results:** We developed the breast cancer epigenomics track hub (BC hub), a resource intended to make it easy for occasional users to find, access and view data of their interest. The BC hub harbors ChIP-seq, ATAC-seq and copy number data from breast tumors, normal breast cells, patient-derived xenografts and breast cancer cell lines in a genome browsable track format. The tracks can be accessed via hyperlinks that automatically configure customized views for different interest groups. Here, we present a detailed description of the resource and informative use cases illustrating its potential in answering specific biological questions.

**Conclusions:** We show that track hubs constitute a powerful way of bringing epigenomics data to the user who could benefit from them. The examples presented highlight the added-value of joint visualization of breast cancer data from different sources. The proof-of-concept provided here exemplifies and underscores the importance of efforts to make biological data FAIR (findable, accessible, interoperable and reusable), and may serve as an encouragement of similar bottom-up initiatives in other research fields. The BC hub is freely accessible at https://bchub.epfl.ch.

## Introduction

### About breast cancer epigenomics data

The epigenome represents the regulatory state of a cell. Sequencing-based epigenomics assays are applicable not only to immortalized cell lines, but also to patient-derived tumor tissues and cells. As such, they constitute a promising and powerful approach to study the diversity of human tumors. Unlike transcriptomics-based characterization of tumor diversity, epigenomic variation between tumors potentially provides mechanistic insights into regulatory defects that operate upstream of gene transcription. Epigenetic assays that have successfully been applied to patient samples include ChIP-seq, ATAC-seq and various types of DNA methylation profiling assays. Pioneering work has been done in the breast cancer field. A brief overview of landmark studies follows.

ChIP-seq was first applied to patient-derived tissue to study estrogen receptor α (ER) and FOXA1 binding in ER+ tumors and metastases from breast cancer patients with good and bad prognosis (Ross-Innes et al., 2012). Subsequently, ER and progesterone receptor (PR) binding sites were mapped in cell line-derived xenografts (Mohammed et al., 2015) and patient-derived xenografts (PDXs) (Severson et al., 2018). The latter study, which focused on male breast cancer, also involved ChIP-seq experiments targeted at androgen receptor (AR), glucocorticoid receptor (GR), GATA-3 and the histone mark H3K4me1. An important study on histone modifications in breast epithelial subpopulations was published in (Pellacani et al., 2016). More recently, ChIP-seq assays for ER were successfully applied to normal breast epithelial cells (Chi et al., 2019).

High resolution digital genomic foot-printing using the DNase-seq technology was first applied to tumor cell lines by the ENCODE consortium (Sabo et al., 2004). Recently, ATAC-seq (Buenrostro et al., 2013), an alternative chromatin accessibility assay, has been applied to circulating tumor cells (CTCs) cultured in vitro and CTC-derived metastases obtained after intracardiac inoculation of mice (Klotz et al., 2020).

### Nature and significance of epigenomics data

Omics experiments are very different from traditional biological experiments. The latter are typically carried out to answer one question at a time regarding a particular gene or protein of interest. Omics data provide answers to thousands of such questions at once. In this capacity, they are of encyclopedic value, as exemplified and explicitly advertised by the ENCODE (encyclopedia of DNA elements) project (Davis et al., 2018). Here are examples of questions with relevance to breast cancer, which could potentially be answered by interrogating public epigenomics data.

1. GREB1 (growth regulating estrogen receptor binding 1) is a well-known ER target gene. Where are its control regions located? Which ones of these regions are bound by ER in normal breast cells, in breast tumors or in tumor-derived cell lines?
2. Kallikrein-3 (KLK3), better known as prostate-specific antigen (PSA) is upregulated in both prostate and AR positive breast cancer. Which are the hormones and hormone receptors responsible for its upregulation in breast tumors?
3. Copy number variation plays an important role in breast cancer. A researcher may be interested in knowing which genomic regions are amplified in the cancer cell line used as model system.
4. Overexpression of AIM2 (absent in melanoma 2) was found to suppress cell proliferation and tumor growth in breast cancer. The expression of this gene is controlled by an enhancer containing binding sites to STAT proteins. Is this enhancer activated by the JAK-STAT signaling pathway in breast cancer?

To enable users to access relevant data and answer such questions more quicky, we developed the breast cancer epigenomics track hub (BC hub). In the following, the above questions will serve as “use cases” for describing this resource.

### Making Epigenomics data FAIR

The scope of the BC hub fits into the broader context of making omics data FAIR: Findable, Accessible, Interoperable and Reusable. Specifically, it makes data reusable by making them instantly accessible and viewable via a few mouse clicks.

The BC hub offers epigenomics data as genome browser viewable tracks. This seems to us the most natural way of bringing the data to the user, as molecular biologists are used to interpret pictorial data presented in papers. In fact, ChIP-seq peaks displayed in dense mode in a genome browser window look like bands on an acrylamide gel.

Free access to epigenomics data is not a problem as such. A wealth of sequencing-based data is freely available from public repositories such as GEO (Barrett et al., 2013) and ArrayExpress (Sarkans et al., 2021). However, often samples are only available as raw data files containing sequencing reads. Downloading and mapping the millions of reads from a single experiment to the genome is computationally expensive and time-consuming, and requires expert skills. Some data are available as browser viewable tracks, but not always in an easily findable and instantly viewable form. By collecting, reformatting, and organizing breast cancer relevant epigenomics data in the BC hub, we aim at making such data more findable, accessible and reusable to the breast cancer research community.

There are parallel ongoing efforts, which help make epigenomics FAIR. The list of public track hubs maintained at UCSC (Rosenbloom et al., 2015) and the recently established track hub registry at EBI enhance findability. The latter is part of the FAIRtracks initiative (Gundersen et al., 2021), which in addition proposes metadata representation standards and offers web services for structured queries. Further of note are Remap (Hammal et al., 2022) and GTRD (Kolmykov et al., 2021), two resources which offer uniformly processed ChIP-seq peak lists in browser viewable BED format, including peak lists from breast cancer data. ChIP-Atlas (Zou et al., 2022) is a portal and data-mining suite for exploring ChIP-seq, ATAC-seq and bisulfite-seq data. It requires a local installation of the IGV genome browser for data visualization.

## Overview of the BC hub

### Scope and guiding principles

The BC hub is a bottom-up initiative with a narrowly defined scope. Wherever possible, we rely on already existing solutions. The UCSC genome browser serves as interactive data access and visualization platform for the user. For upstream data processing and track generation, we rely on protocols previously developed for other resources, namely the mass genome annotation (MGA) data repository (Dreos et al., 2017) and the eukaryotic promoter database EPD (Meylan et al., 2020). For quality control, we use programs from the ChIP-Seq tools (Ambrosini et al., 2016).

We will briefly introduce the BC hub from a user perspective with an example. First, we present a genome browser view and explain what the displayed tracks represent and how they could be interpreted. Next, we describe how the users generate the view via the BC hub.

### Exploring epigenomics data with the BC hub

Fig. 1 presents a genome browser screenshot showing breast cancer epigenomics data for the upstream region of the GREB1 gene, a known target of ER. ChIP-seq data from four studies are shown, covering various types of cellular sources: cancer cell lines, surgically removed tumors and metastases, normal breast epithelial cells and PDXs. The tracks are shown in “dense” display mode rather than “full”, which is more commonly seen in research papers. Besides saving space, this has the advantage that binding peaks appear like bands on an acrylamide gel, which may be more intuitive and easily interpretable by the trained eye of a wet lab biologist.

**Figure 1:**
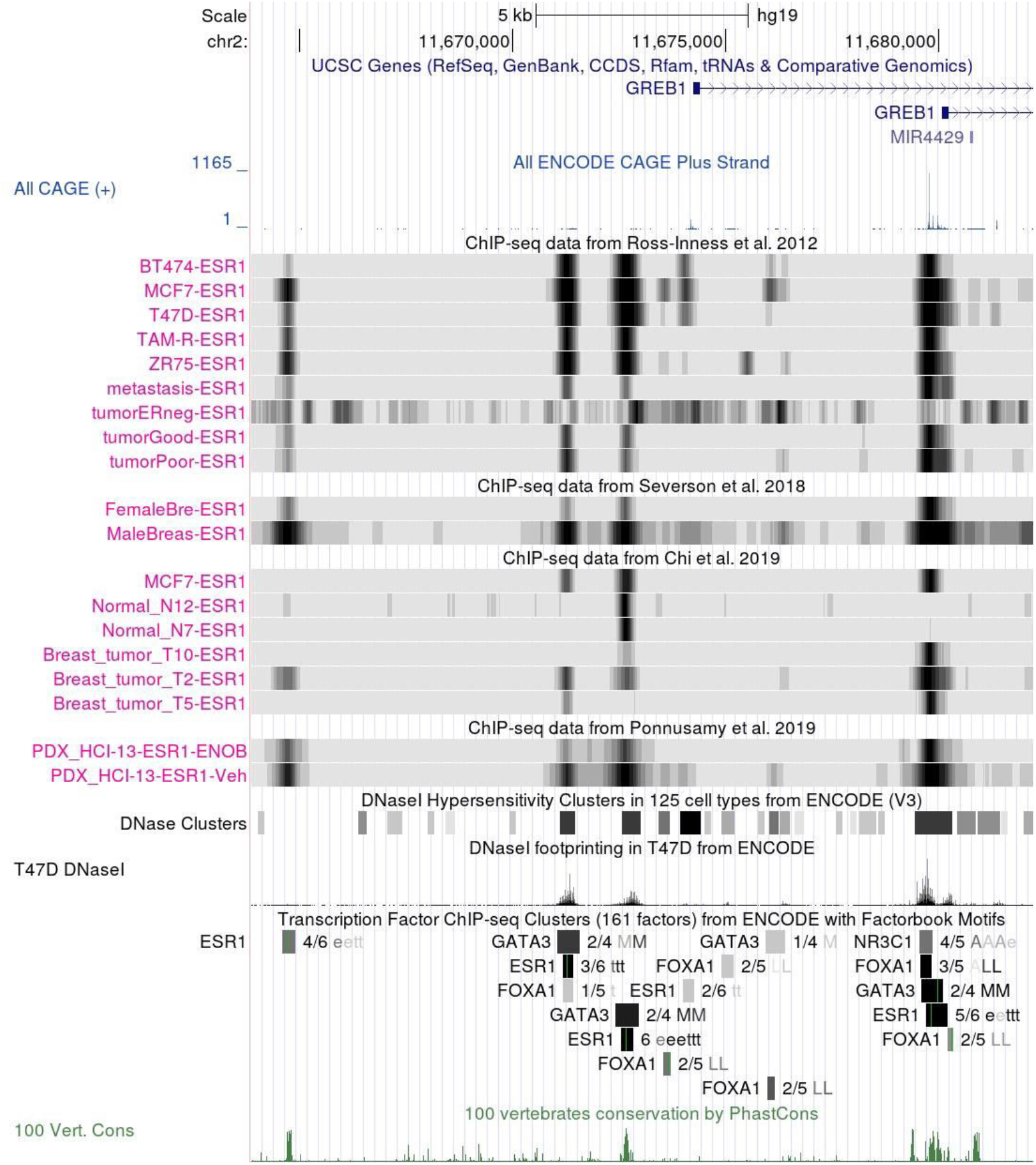
ESR1-centric view of the GREB1 promoter. Top: UCSC Gene tracks showing the 5′-terminal exons of two GREB1 transcripts; ENCODE All CAGE track showing transcription start site usage at single base resolution. Middle part: ER ChIP-seq tracks from the BC hub. Data were generated from a variety of cellular sources: breast cancer cell lines, surgically removed tumors and metastases, normal breast epithelial cells and patient-derived xenograft. Bottom: Additional tracks from the UCSC genome browser database, including chromatin accessibility profiles and merged ChIP-seq peaks from ENCODE plus the PhastCons cross-genome conservation track. A detailed interpretation of this picture is presented in the main text. The BC hub tracks shown are based on data from (Chi et al., 2019; Ponnusamy et al., 2019; Ross-Innes et al., 2012; Severson et al., 2018). Abbreviations in track names: ENOB, enobosarm; Veh, vehicle.

Let’s now have a closer look at these tracks. The browser window shows four major ER (denoted ESR1 in all track names) peaks, which will be referred to as regions 1 to 4 in the sequel. We can see that these regions are differentially occupied in different subsets of samples. Region 4 (the rightmost) corresponds to the main promoter of GREB1 according to the CAGE data track. It is bound by ER in most samples but curiously not in normal breast epithelial cells. On the other hand, regions 1 and 2 are strongly bound in all cell lines, but less frequently and to varying degrees in native tumors and PDXs. Region 3 is the only one bound by ER in normal breast cells.

Also included in the browser screenshot are tracks from other sources, mostly from the UCSC genome browser database (Lee et al., 2022). These tracks are meant to provide contextual genomic information supporting the interpretation of the breast cancer tracks. On top is the UCSC Genes track showing the 5’-termInal exons of two alternative transcripts. Just below is a CAGE track based on data from ENCODE (Djebali et al., 2012) showing the cumulative transcription start site (TSS) usage in all 31 normal and cancer cell lines analyzed. This track is imported from the EPD promoter viewer track hub (Meylan et al., 2020). The CAGE data confirm that the ER bound region 4 corresponds to the major GREB1 promoter.

Just below the breast cancer tracks appear two chromatin accessibility tracks from the ENCODE consortium. The first track shows merged DNaseI hypersensitive regions from 125 cell lines. The grey scale reflects the average intensity of the DNaseI signal. A click on the rectangular icons opens a new window showing the respective signal strength for all cell lines assayed. The second track shows the DNaseI cutting frequency in T47D cells at single-base resolution. We note that region 1 is not reported to be accessible in any of the ENCODE cell lines whereas the other 3 regions all are. The absence of region 1 in the ENCODE DNaseI track may not be totally surprising in view of the overall weaker ER signal intensity seen in the ChIP-seq tracks.

Just below the T47D DNaseI track appears the composite ENCODE ChIP-seq cluster track. Each grey bar represents the merged binding regions for one TF in 91 cell types. A click on the grey bar will open a new window listing the cell lines, in which assays have been performed and binding peaks were found. The thin green vertical bars mark computationally predicted binding motifs defined by position specific scoring matrices from Factorbook (Wang et al., 2013). The complete track provides data for 161 different TFs, which can individually be selected for display. Not surprisingly, the four ER binding regions found in tumor tissue are confirmed by the ENCODE ChIP-seq data. All four regions contain ER motifs suggesting that ER binds directly to the DNA rather than by piggy-packing on another TF. Note further that the weakly bound region 1 is the only one, which is not simultaneously bound by any of the three other TFs selected for display.

The bottom-most PhastCons (Siepel et al., 2005) track indicates cross-species sequence conservation. We note that region 1, though largely inaccessible according to chromatin assays and not bound by ER in normal breast cells, is nevertheless highly conserved. This suggests that this region has a vital function in the regulation of GREB1 expression in normal tissues, though perhaps not in the breast. Conversely, the lack of cross-species conservation of region 2 suggest that this region is accidentally activated in breast tumors and may be causally involved in tumorigenesis.

### Accessing the BC hub

The BC hub not only offers epigenomics data as browser viewable tracks, it also provides mechanisms designed to find, access and combine data from all over the world in a single genome browser window. The BC hub home page features so called session links to this end. A mouse-click on a session link connects the user to the UCSC genome browser host and sends back a customized view. For instance, the customized view shown in Fig. 1 has been generated with a link to an ER-centric view for GREB1. Accessing the genome browser via a session link triggers several actions on the browser host: connecting the new browser session to the BC and other public hubs, selecting a track subset for visual display, defining the order of the tracks and setting their display mode. The initial view typically serves as a starting point for further customization. To obtain the compact view in Fig. 1, many initially visible tracks were hidden, and the preselected subset of breast cancer relevant TFs from the ENCODE ChIP-seq cluster track was further reduced to the following four factors (identified by gene symbol): ESR1 (ER), NR3C1 (GR), FOXA1 and GATA3. Conversely, the initially hidden “all ENCODE CAGE” track from the EPD viewer hub was turned on in full display mode. These changes were interactively applied via the track set selection and configuration menus provided in the lower part of the genome browser window.

## Implementation

### Overview of the content generation process

The BC hub physically consists of preprocessed public data in a genome browser track format (bigWig) plus additional files, which enhance accessibility, findability and usability of the data. The latter include the track database and the already mentioned session files.

The dataflow during the import and data preprocessing steps is schematized on the left side of Fig. 2. The source data from different repositories offered in different formats are imported into the MGA repository, which serves as an infrastructure for uniform data preprocessing, quality control and metadata curation (Dreos et al., 2017). (The MGA repository already serves this purpose for the EPD promoter viewer hub.) All data in the MGA repository are represented in SGA (simple genome annotation) format, the working format of the ChIP-Seq tools (Ambrosini et al., 2016). From there, browser viewable tracks in bigWig format are generated using the data processing pipelines of EPD. The only exception to this rule is the procedure for generating the copy number variation (CNV) tracks (see Methods).

**Figure 2.**
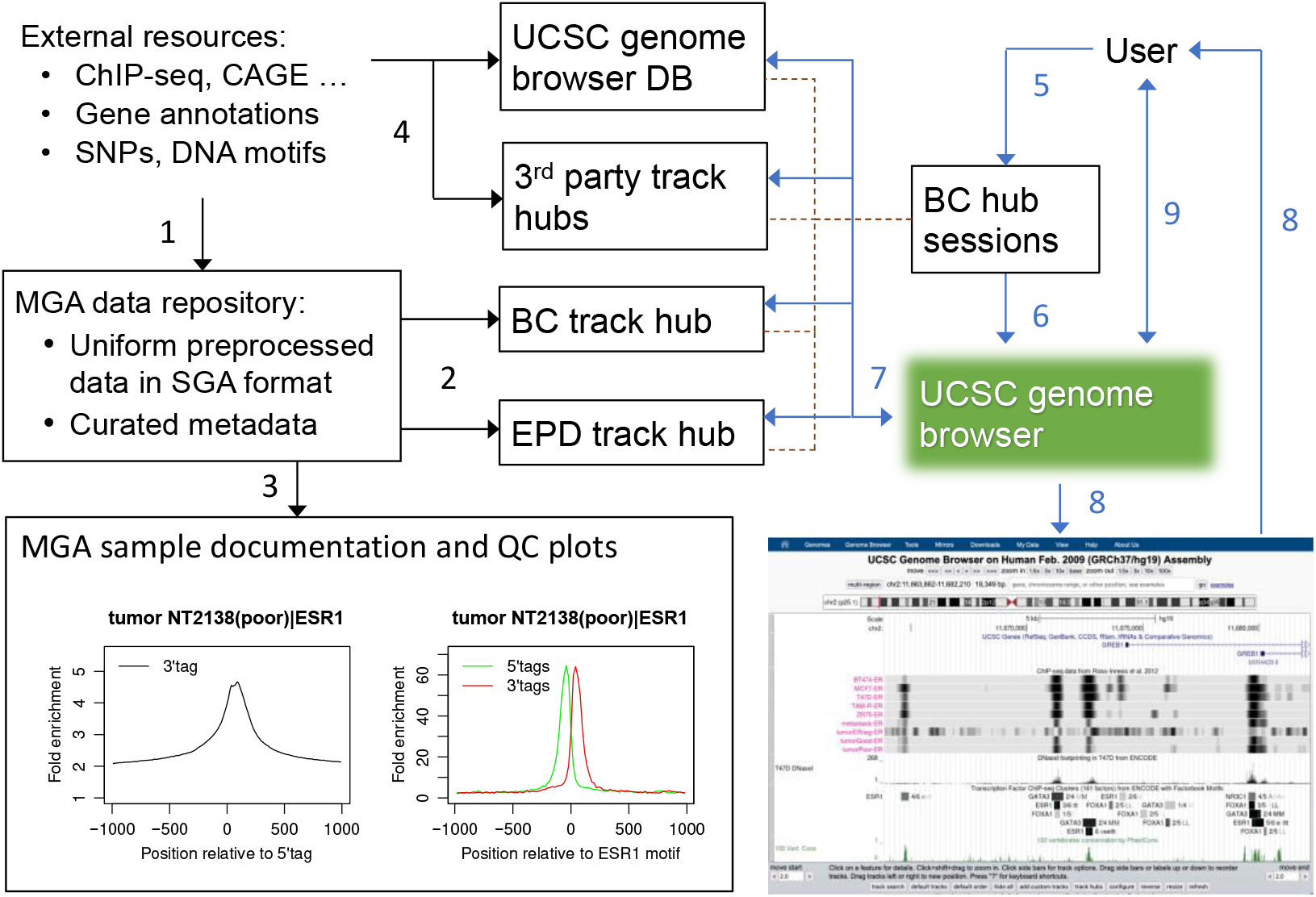
Data Flow during backend preprocessing and interactive data access. 1) Reformatting and import of public data into the MGA repository. 2) Reprocessing and export of selected data sets to the breast cancer and EPD track hubs. 3) Generation of textual documentation and QC plots for the breast cancer samples; these documents remain in the MGA domain. 4) Outside BC hub: public data are preprocessed and incorporated into the UCSC genome browser database and third-party track hubs. 5) Users access the BC hub via a link to a session posted on the BC hub home page; session files reside on the web server hosting the BC hub. 6) The UCSC genome browser receives instructions from a session file as to how to configure a customized view. 7) The genome browser extracts the required track segments from its own database and remote track hubs. 8) The genome browser composes the customized view and returns it to the user. 9) Later during the session, the user may interactively change the view; each such interaction may trigger new data transfers between the genome browser host and remote track hubs. Color codes: black arrows indicate data flow during preprocessing; blue arrows indicate data transfers during an interactive browser session; broken lines represent cross-references from the session file to tracks in the genome browser database or external data hubs.

The BC hub further relies on the MGA repository for track annotation. The integration of an epigenomics data series into the MGA repository includes data curation efforts. Biologically relevant sample information is first automatically imported from machine-readable metadata files provided by the source repositories. This information is subsequently verified, complemented and if necessary corrected upon critical reading of the corresponding scientific articles by human curators. Comprehensive sample annotation for each series is posted on the MGA web server along with quality control plots.

### The track database

The track database (track DB) is a computer readable text file providing browser actionable information about all tracks in the hub. It is publicly accessible at: https://ccg.epfl.ch/ucsc_hubs/breastca/hg19/trackDb.txt. One of the main functions of the track DB is to group tracks into track sets (called composite tracks inside the track DB file). In the BC hub and MGA repository, we usually mirror the organization of samples in the source repositories, GEO (Barrett et al., 2013) or ArrayExpress (Sarkans et al., 2021). The currently available track sets are listed in Table 1. Each set corresponds to one publication. The track DB further defines the visibility status of each track (shown or hidden), the order in which the tracks appear in the browser window and various display options (color, track height, etc.). All these settings can be changed by the user at any time during an interactive browser session. They can also be over-written by session files (see below).

**Table 1.** Current track sets of the BC hub.

| UCSC track name | Track description | Track resolution [bp] | Original dataset | GEO | Tissue | Tumor Grade | Condition | MGA series name | PMID |
| --- | --- | --- | --- | --- | --- | --- | --- | --- | --- |
| 1 Siersbaek 2020 | ESR1, FOXA1, STAT3, and H3K27ac ChIP-seq | 50 | PRJNA518884 | GSE126006 | Primary ER+ breast tumors, MCF7 and T47D | n/a | Vehicle and IL6-treated | siersbaek20 | 32679107 |
| 2 Klotz 2020 | ATAC-seq | 1 | SRP139038 | GSE112852 | Breast cancer CTC lines and CTC-derived metastases in mouse | n/a | n/a | klotz20 | 31601552 |
| 3 Ponnusamy 2019 | ESR1, AR and FOXA1 ChIP-seq | 50 | SRP187889 | GSE128018 | Breast cancer xenografts | n/a | Vehicle and enobosarm-treated | ponnusamy19 | 31698248 |
| 4 Nacht 2019 | PGR, CEBPA, RAD21 and TOP2A ChIP-seq | 50 | SRP201251 | GSE132649 | T47D | n/a | Progestin (R5020) treated | nacht19 | 31373033 |
| 5 Chi 2019 | ESR1, GRHL2 and H3K27ac ChIP-seq | 50 | SRP108616 | GSE99680 | Normal breast cells, tumors and MCF7 | n/a | n/a | chi19 | 31110002 |
| TRACK SUBSET: ChIP-seq and CNV |  |  |  |  |  |  |  |  |  |
| 6 Severson 2018 | ESR1, AR, PGR, NR3C1, FOXA1, GATA3, and H3K4me1 ChIP-seq | 50 | SRP119087 | GSE104399 | Male and female breast tumors, and MCF7 | n/a | n/a | severson18 | 29396493 |
| 7 Severson 2018 | Genome-wide log2-transformed, normalized (ratios of) read counts (CopywriteR from Bioconductor) | 50 | SRP119087 | GSE104399 | Male and female breast tumors, and MCF7 | n/a | n/a | severson18 | 29396493 |
| TRACK SUBSET: ChIP-seq and ATAC-seq |  |  |  |  |  |  |  |  |  |
| 8 Takaku 2018 | GATA3 and mutated (ZnFn2) GATA3, ESR1, PGR, FOXA1, and PolII ChIP-seq | 50 | SRP108337 | GSE99479 | T47D and GATA3 mutant cell line (CR3) | n/a | Vehicle and P4 | takaku18 | 29535312 |
| 9 Takaku 2018 | ATAC-seq | 1 | SRP108337 | GSE99479 | T47D and GATA3 mutant cell line (CR3) | n/a | n/a | takaku18 | 29535312 |
| 10 Maturi 2018 | ZEB1 ChIP-seq | 50 | E-MTAB-5241 | n/a | Triple-negative breast cancer cell line Hs578T | n/a | n/a | maturi18 | 29744893 |
| 11 Singhal 2016 | ESR1, PGR, and p300 ChIP-seq | 50 | PRJNA318944 | GSE80367 | T47D | n/a | Hormone treated (E2, R5020) | singhal16 | 27386569 |
| 12 Mohammed 2015 | ESR1, PGR, and p300 ChIP-seq | 50 | SRP057755<br>SRP057756<br>SRP057757 | GSE68359 | MCF7, T47D, and xenografted MCF7 | n/a | Estrogen-treated and with/without progestins (Progesterone/R5020) | mohammed15 | 26153859 |
| 13 Ross-Inness 2012 | ESR1 and FOXA1 ChIP-seq | 50 | SRP032421 | GSE32222 | Primary ER-positive breast tumors from patients, distant ER-positive metastases, and cancer cell lines (MCF7, T47D, ZR75, BT474) | n/a | n/a | ross12 | 22217937 |

The track DB not only determines the default display of the tacks in the graphical genome view but also the organization of the track configuration menus in the lower part of the genome browser window. The entire BC hub appears as one section under the title “Breast Cancer Epigenomics Track Hub” (Fig. 3A). Each track set can be accessed via a pull-down menu and a hyperlink. The pull-down menu serves to change the visibility status of the entire track set. The hyperlink takes the user to a detailed track selection and configuration page (Fig. 3B). The track DB format offers the possibility to assign attributes (terms from a controlled vocabulary) to each track in a track set. In the example shown, the tracks have been grouped by tissue source and ChIP-seq target. This results in the display of a table that enables user to select subsets of tracks by a single click on a checkbox. Alternatively, multiple choices from a long list of attributes can be offered by a pull-down menu (Fig. 3C), exemplified by the ENCODE ChIP-seq cluster track from the UCSC genome browser database.

#### Session links

Public track hubs can be accessed manually during a browser session or via a session link. The session link is a composite URL including the URL of a session file as a parameter. A session file stores the track selection and display settings of a current browser window. It can be saved by the user to a local disk at any time during a session. Later, it can be uploaded to the genome browser in order to reproduce exactly the same view. On the BC hub home page, we offer session links generated in this way to customized views for different user communities or thematically defined focus areas. The session mechanism is also used to take users directly to genes of interest such as GREB1 (Fig. 1). Session files not only over-write default settings in the track DB. They also establish connections to other public track hubs. For instance, the session link used as starting point to generate Fig. 1 automatically connects the user to the EPD promoter viewer hub and a public Cancer Genomics hub harboring tracks from TCGA and ICGC.

Note that the session links of the BC hub guide users from a specialized community to “needles in a haystack”: a tiny subset of interesting tracks from an ever-growing number of publicly available tracks. In doing so, they help make epigenomics data FAIR, in particular findable.

#### Data flow during a UCSC genome browser session

The UCSC genome browser is a data visualization platform, supporting on-the-fly data integration and visualization. The right side of Fig. 2 illustrates the data flow during an interactive browser session. Users of the BC hub access the UCSC genome browser host via a session link posted on the BC hub home page. A mouse-click on the link sends a session file to the browser host. The session file contains instructions to compose a customized view for the user. Upon receipt, the browser host connects to external hubs, extracts data from selected tracks for the requested genomic region, adds additional tracks from the server-resident UCSC genome browser data base to the view, configures the display of all selected tracks according to the instructions in the session file, and finally sends back an image to the viewing device (desktop, laptop or smartphone) of the users. Once connected to the browser, the user can further modify the display settings, hide or add new tracks, or navigate to another genomic regions via the menu buttons posted in the browser window beneath the genome view. Each such actions triggers new data flows of the same kind as described above.

It is important to recognize that efficient data transfer protocols leveraging indexed “big data” formats such as bigWig (Kent et al., 2010) play a mission critical role in decentralized data integration via a track hub. Only the track DB file needs to be stored at the browser host when an external hub is connected to a session. The voluminous track files remain at the hub host. Later, only tiny portions of these files are temporarily transferred to the browser host upon selection of a new genomic region by the user. In retrospective, the introduction of big data formats has been a real game-changer in genomics data visualization.

#### Track documentation

For each track set, there is a short documentation page structured according to UCSC guidelines, which can be accessed via the link provided in the BC hub track selection menu (Fig. 3A). This page contains hyperlinks to the more comprehensive documentation pages and quality-control (QC) plots provided by the MGA repository.

**Figure 3.**
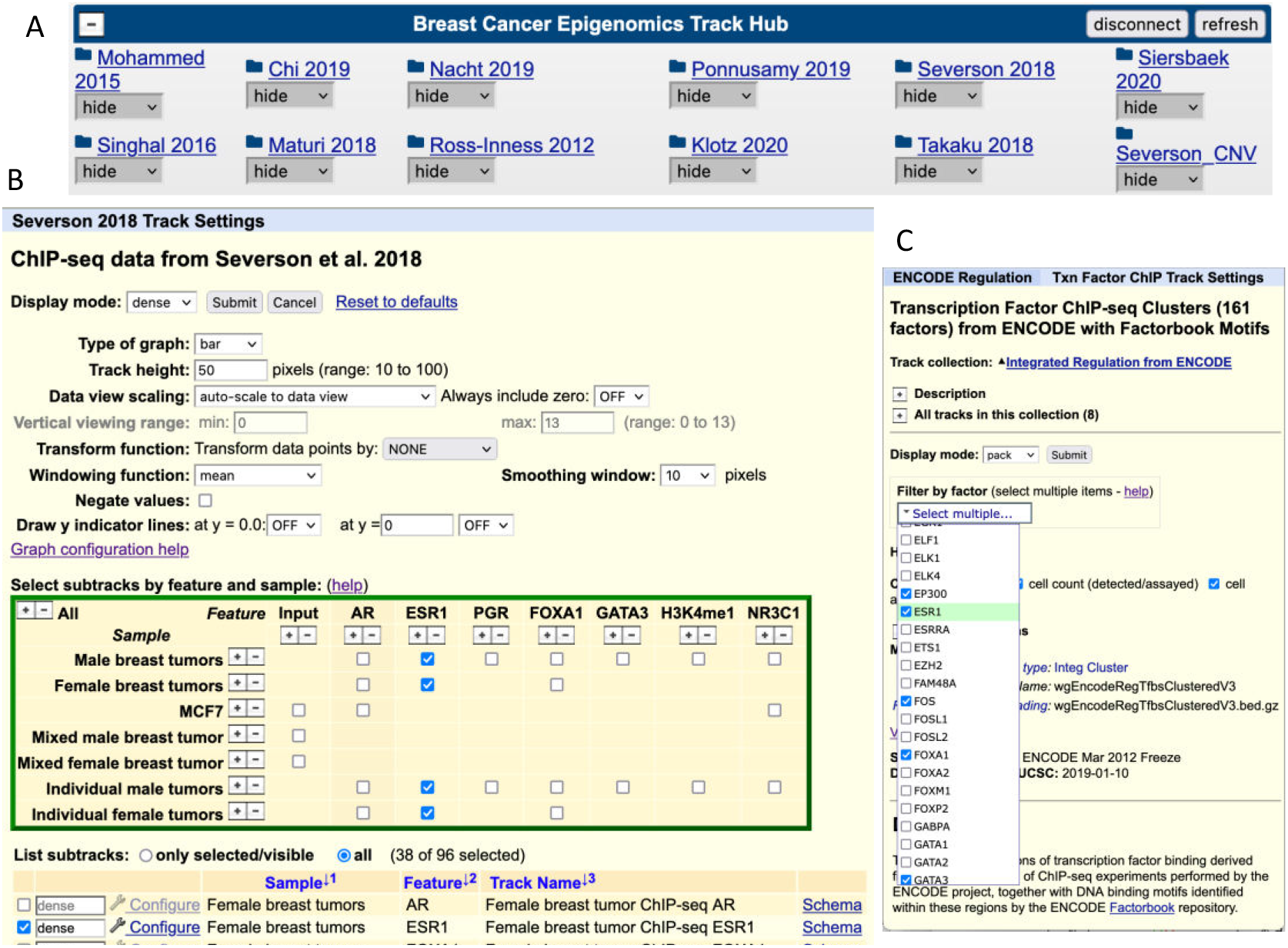
Track selection menus. A. Track sets from the BC hub appearing in the UCSC genome browser track selection and configuration menu. The pull-down menus allow users to directly change the display mode of the visible tracks. Clicking on the track set title takes the user to the track set configuration menu. **B**. Configuration menu for the Severson et al. 2018 track set. The table allows users to select and an unselect groups of related tracks at once based on track attributes (sample types or ChIP-seq target) defined in the track hub database. **C**. Pull-down menu for selecting TFs from the ENCODE ChIP-seq cluster track exemplifying an alternative mechanism to select object types from a long list.

Concise information about individual tracks is communicated to the users via three descriptive elements: (i) the sample description included in the MGA series documentation page, (ii) the track name and (iii) track description. The latter two elements are stored in the track DB of the BC hub. An example is presented below. The track chosen (visible in Fig. 1) is from the “Ponnusamy et al. 2019” track set.

MGA sample description: BC PDX HCI-13|AR|ENOB|rep1
Track name: PDX_HCI-09-AR-ENOB
Track description: PDX HCI-09 ChIP-seq AR, ENOB

Note that the MGA sample description is structured. It consists of four fields separated by vertical bar. The exact usage of the four fields varies somewhat depending on the assay type. For ChIP-seq, they correspond to cellular source, antibody target, treatment, and replicate identifier. The series documentation page provides complementary information such as explanations of abbreviations, to help users decode the concise sample descriptions (see https://ccg.epfl.ch/mga/hg19/ponnusamy19/ponnusamy19.html, with regard to the above example).

The MGA series documentation page is meant to be directly viewable by the users, unlike the track name and descriptions, which are displayed through the genome browser software. The names appear on the left side of the corresponding tracks (Fig.1), the descriptions on top of the tracks if the display mode is set to full instead of dense. The display mode can be changed via the track set configuration menu (Fig. 3B).

To achieve uniformity across the BC hub, we try to adhere to some nomenclature standards. In the absence of such standardization in the original data sources, this is a delicate a task that has to be tackled with caution. Changing names and abbreviations of cell lines, assays or proteins could make it more difficult for the user to match track names seen in the browser window to samples referred to in the corresponding publication. The goal of any such effort must therefore be to find an optimal compromise between closeness to the sources and uniformity within the hub.

Currently, we only try to standardize the names of the proteins targeted by ChIP-seq experiments, for which we use HGNC gene symbols (Tweedie et al., 2021). This ensures consistency with track sets from other sources included in the customized views. For instance, estrogen receptor α (ER) appears as ESR1 in the track set selection table from the BC hub (Fig. 3B), as well as in the pull-down menu for the ENCODE ChIP-seq clusters track (Fig. 3C). Usage of gene symbols is also recommended by the FAIRtracks initiative and commonly applied in other genome informatics resources.

#### Quality control

Visual exploration of tracks by a trained eye often provides a good intuitive judgement of the data quality and exploitability (Fig. 4A). Nevertheless, intuition can be misleading, and a small genomic region can never be representative of the genome as a whole. Therefore, we provide two types of global QC plots for a subset of tracks: ENCODE-style strand cross-correlation graphics for all ChIP-seq samples and signal-enrichment plots for TFs with a known binding motif (Fig. 2). The latter are aggregation plots (Jee et al., 2011) showing the density of 5’ and 3’ tags (reads mapped to the + and – strand of the genome) around the 1000 computationally identified binding motifs with the highest read coverage.

**Figure 4.**
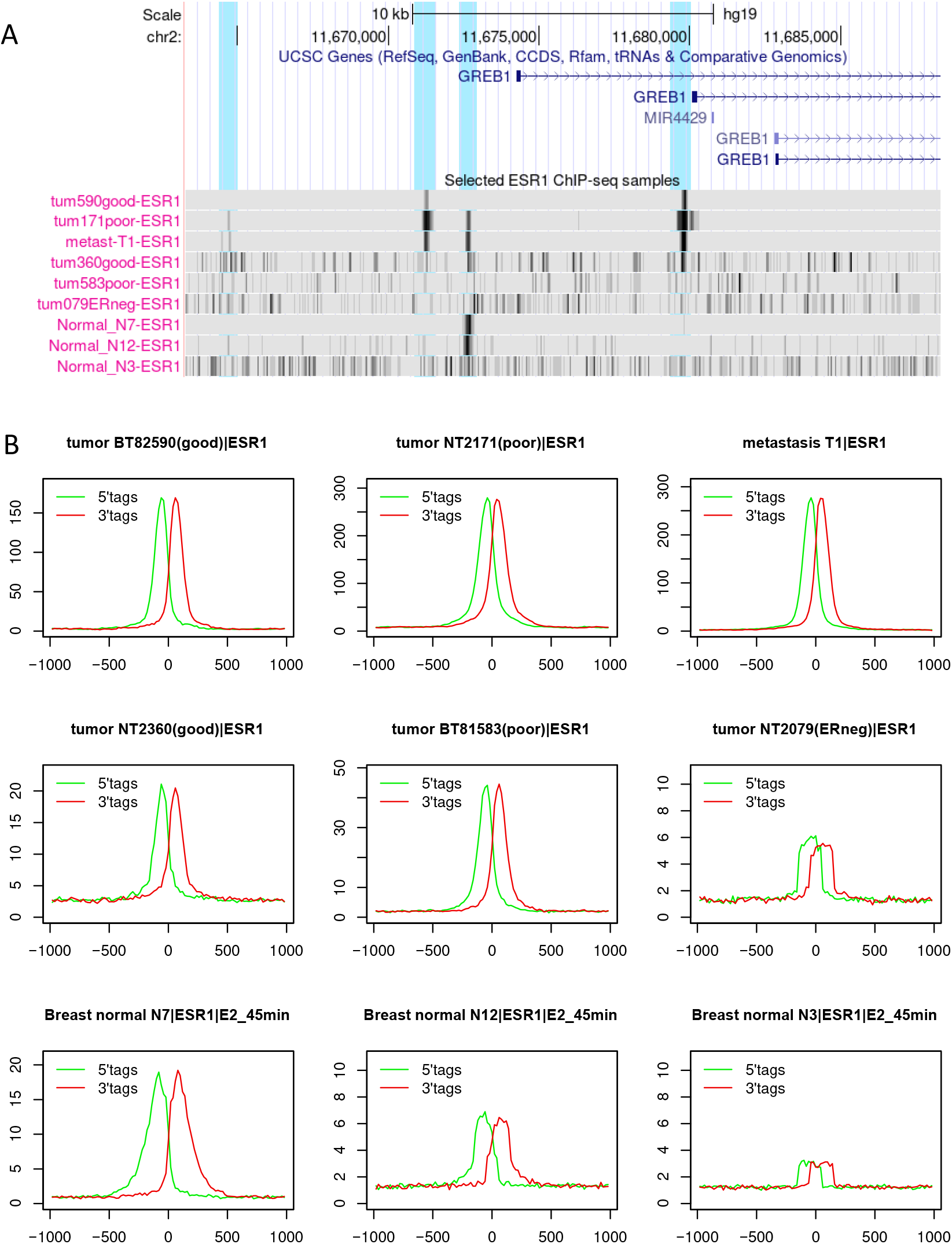
Quality control information for selected ChIP-seq samples. A Local view: genome browser screenshots showing selected examples of ChIP-seq tracks for tumor samples and normal breast epithelial cells in a narrow genomic region. **B** Genome-wide analysis: Signal enrichment plots around ER binding motifs with high read coverage. The green and red curve represent the density of reads mapping to the + and – strand of the genome, respectively (see Methods for details). The vertical axis reflects fold change over genome average. A detailed interpretation of the tracks and plots is presented in the main text. The BC hub tracks are based on data from (Chi et al., 2019; Ross-Innes et al., 2012).

ChIP-seq data from patient tissue can be of highly variable quality. This is illustrated with selected breast tumor and normal breast tissue samples in Fig. 4. A local view of a narrow genomic region (same as in Fig. 1) is provided in Fig 4A. Note that the signal intensity is displayed in dense mode and scaled to the maximum viewing range. With these settings, strong narrow bands on a clean background are indicative of high data quality. Conversely, background noise over the entire viewing range could either be due to low data quality or the absence of true binding peaks in the selected viewing range.

Fig. 4B presents global (genome-wide) signal enrichment plots. Note the expected shift of about 100-150 bp between the 5’ and 3’ tag distributions. Data quality is reflected by the height of the peaks and smoothness of the curves. Overall, we see good agreement between the noisiness of the genome browser tracks and data quality suggested by the signal enrichment plots. However, there are some notable exceptions. The tumor samples 360good and 583poor (good and poor relate to prognosis inferred from biomarkers) look much better in the QC plots than in the genome browser. The opposite is true for the normal breast tissue samples N7 and N12. Consistently low data quality is indicated, as expected, for the ER negative tumor sample but (disappointingly) also for the normal sample N3.

#### User support and Documentation

We currently offer user support by two means, a beginner’s tutorial and a session gallery.

The tutorial consists of a guided tour for new users. During the tour, participants will learn how to access and visualize data, how to reconfigure the track sets and how to add new tracks to the view and hide others. Users will also learn to upload new tracks via URL or from disk, possibly their own data. The document further features links to tutorials and documentation pages of the UCSC genome browser.

The session gallery consists of a series of browser screenshots related to use cases. An accompanying text explains the biological background and walks the beholder through the picture. The idea to teach users in this way how to explore public data was inspired by the ENCODE session gallery posted on the UCSC genome browser website. Our session gallery is primarily intended to be an interpretation guide for BC hub users. However, it could also serve as an E-learning tool that introduces students to epigenomics and high-throughput epigenomics assays by examples.

### Use cases

In this Section, we will illustrate the encyclopedic value of epigenomics data and the power of the BC hub in leveraging such data, with three examples taken from the session gallery. These use-cases correspond to questions raised in the introduction.

#### Androgen and progesterone receptor binding at the KLK3 locus

Kallikrein 3 (KLK3), better known as prostate-specific antigen (PSA), is also expressed in the breast and has been proposed as a biomarker for breast cancer (Hanamura et al., 2019). In breast cancer cell lines, the gene appears to be under the control of both androgen and progesterone (Paliouras and Diamandis, 2007). It is therefore not clear which one of the two hormones drives KLK3 expression in breast tumors. Answering this question is further complicated by the fact that the corresponding receptors recognize virtually identical sequence motifs. Interrogating public ChIP-seq data for PR and AR in patient-derived tumors and tumor cell lines may help resolve this issue.

**Figure 5:**
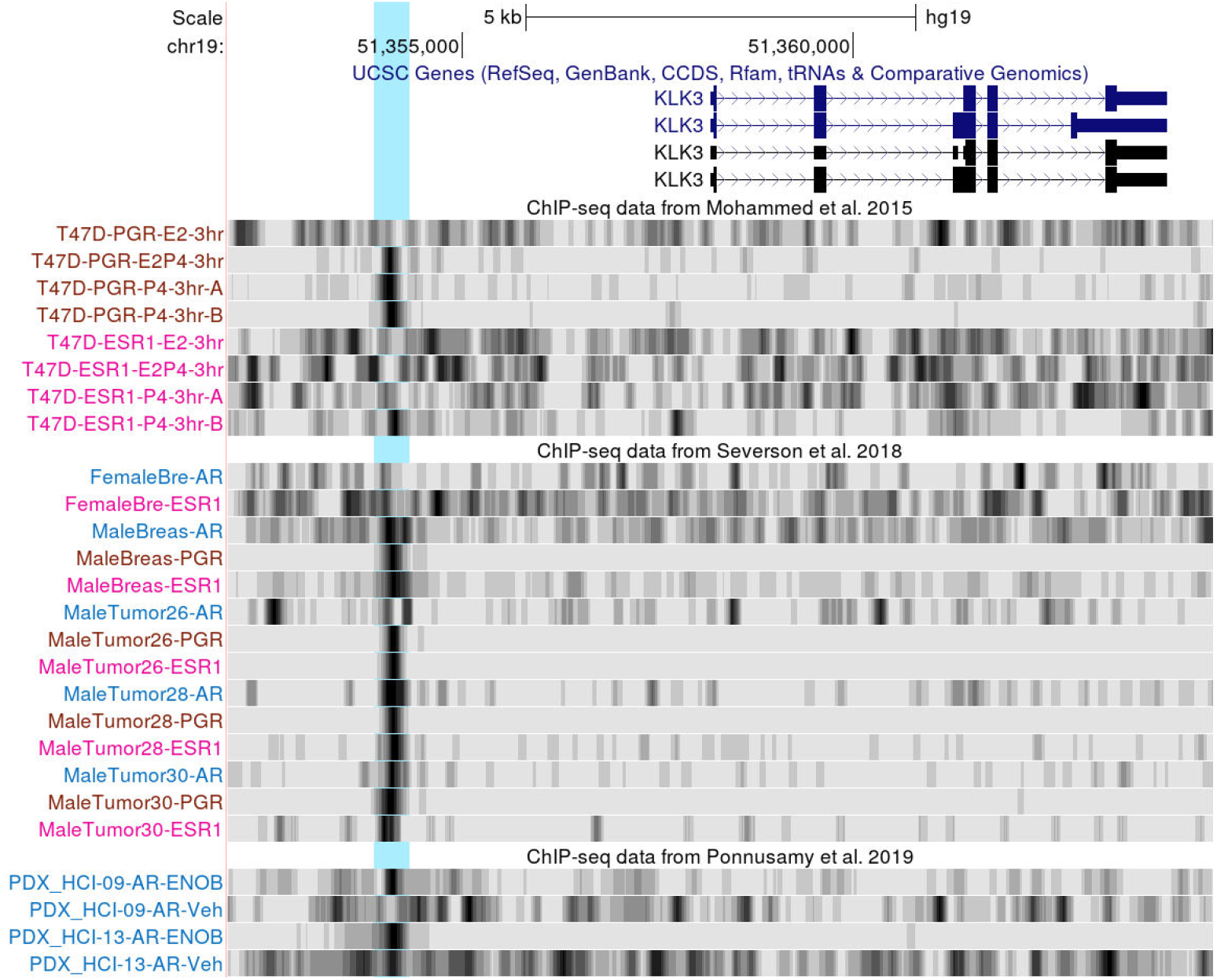
Androgen and progesterone receptor binding at the KLK3 locus. The vertical pale blue bar marks an androgen-responsive enhancer derived from an LTR40a transposable element. BC hub track sets from top to bottom: (i) ChIP-seq data for ER and PR in T47D cells under different treatment conditions; (ii) ChIP-seq data for hormone receptors from pooled and individual female and male breast tumor tissue; (iii) ChIP-seq data for AR from female breast tumor-derived xenografts with and without androgen agonist treatment. All tracks are displayed in “autoscaled to data view” mode (Fig. 3B). Selective binding to the highlighted enhancer regions must therefore be judged based on the signal-to-noise ratio. A detailed interpretation of this picture is presented in the main text. The BC hub tracks shown are based on data from (Mohammed et al., 2015; Ponnusamy et al., 2019; Severson et al., 2018). Abbreviations in track names: E2, estradiol; P4, progesterone; ENOB, enobosarm; Veh, vehicle.

Fig. 5 shows a PR/AR-centric view of the KLK3 locus. Highlighted in orange is a known androgen-responsive enhancer derived from a retro-transposed LTR40a element. This enhancer is believed to be responsible for the up-regulation of KLK3 in LNCaP prostate cancer cells. The ChIP-seq tracks from Mohammed et al. 2015 indicate that the enhancer region is indeed bound by PR (denoted PGR in the track names) in T47D cells, but only upon progesterone treatment. Estrogen alone does not induce binding of PR, and neither hormone induces ER binding to this region. In three different male breast tumors (tracks from Severson et al. 2018), the enhancer is consistently bound by PR and ER, but only in two of them also by AR. Unfortunately, there are no PR binding data for female breast cancers. However, the only female tumor, which has been assayed for AR binding, doesn’t appear to be bound by AR.

The overall picture is completed by AR tracks for mouse xenografts derived from female breast tumors. There, the enhancer is clearly bound by AR upon treatment with the androgen receptor agonist enobosarm. In summary, the ChIP-seq tracks in Fig. 5 show that the highlighted enhancer can be bound by PR and AR in breast tumors, and that receptor binding can be induced in cell lines by either hormone. However, the identity of the hormones binding to the receptors and the respective downstream effects of PR versus AR binding require further investigation.

### Copy number variation in the 17q23 chromosomal region

Segments of the 17q23 chromosomal region are frequently amplified in breast tumors and breast cancer-derived cell lines, including MCF7 (Sinclair et al., 2003). It has repeatedly been observed that copy number variation can be inferred from sequencing-based epigenomics data, including ChIP-seq. Amplified regions are often visible in native ChIP-seq tracks at low resolution display. More sophisticated protocols exist to infer quantitative estimates of local copy number variation. This could be important for elucidating the causes of altered expression of cancer-relevant genes in individual tumors. Two alternative hypotheses have to be considered. The change in expression could merely reflect the change in copy number, or it could be the consequence of regulatory defects acting upstream of transcription. We use the 17q23 region as an example to illustrate visualization of copy number variation using ChIP-seq data from two studies (Ross-Innes et al., 2012; Severson et al., 2018).

The browser screenshot in Fig. 6 covers a large (~7 MB) genomic region. Shown are native ChIP-seq tracks for ER and FOXA1 along with computationally inferred copy number tracks. At this resolution, the genome amplification reported in MCF7 cells is clearly visible at the center of cytoband 17q23.2 (tracks from Ross-Innes et al. 2012). A recently published method called CopywriteR (Kuilman et al., 2015) was then used to generate computationally inferred copy number tracks from raw ChIP-seq counts (Copy number tracks from Severson et a. 2012). This method discards reads falling into peak regions and applies a base-composition correction to the read densities outside peak regions (Kuilman et al., 2015). Strong amplicons are often already visible in native ChIP-seq data. We notice, for instance, that the native ChIP-seq tracks for ER and FOXA1 in MCF7, and the computationally derived copy number track for MCF7 are virtually indistinguishable.

**Figure 6.**
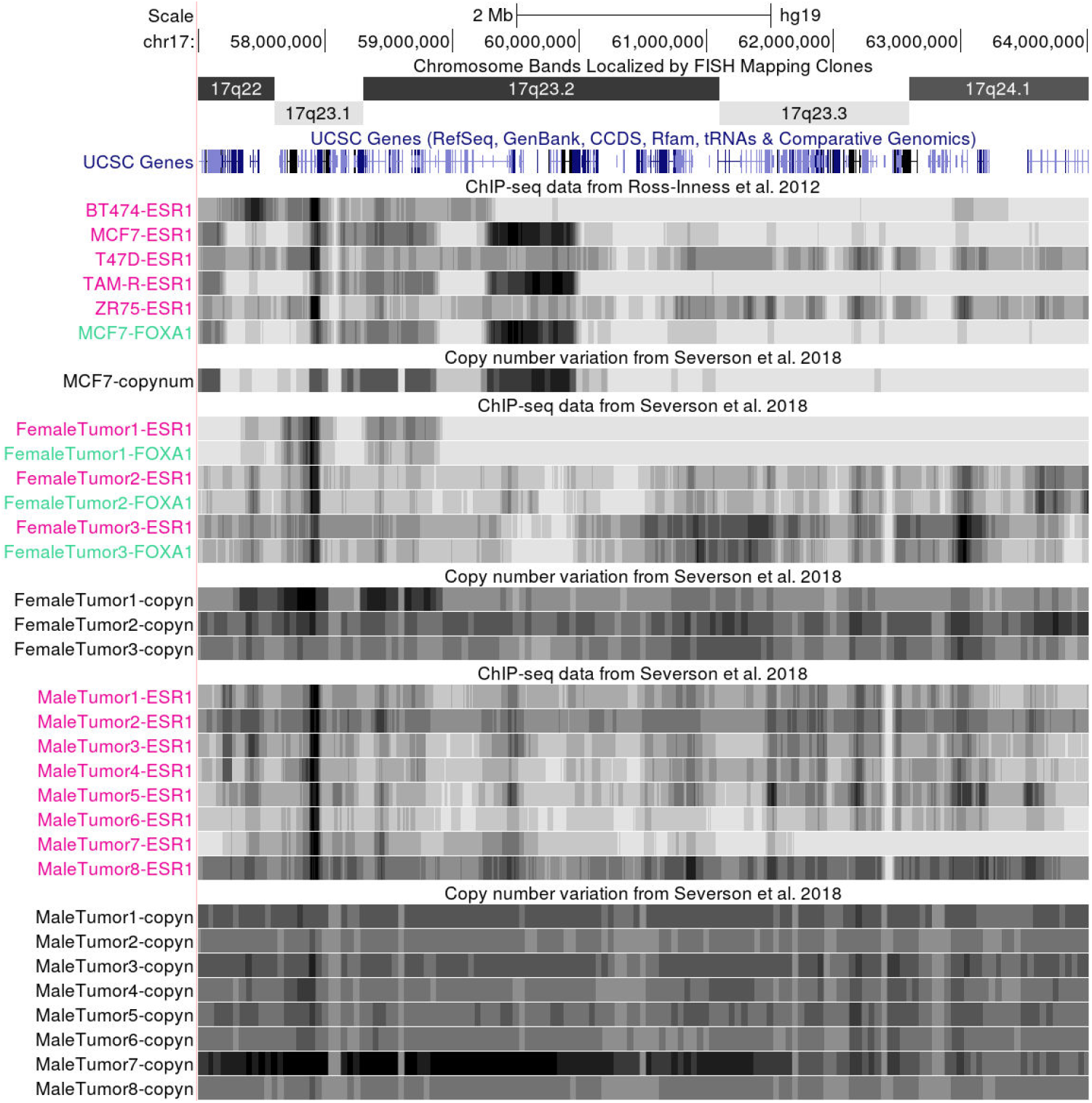
Copy number variation revealed by ChIP-seq at 17q23. This browser screenshot covers a large (~7 MB) genomic region including the 17q23 amplicon frequently present in breast tumors and cell lines, including MCF7 (Liu et al., 2018; Sinclair et al., 2003). Unmodified ChIP-seq tracks are compared to computationally inferred coy-number tracks generated with the CopywriteR software (Kuilman et al., 2015). Tracks sets from top to bottom: (i) Cytobands and dense gene display from the UCSC genome browser database. (ii) Various ChIP-seq tracks from cell lines and inferred copy number track for MCF7. Note the strong congruence of the three MCF7 tracks (ESR1, FOXA1 and copynum). (iii) ChIP-seq and inferred copy number tracks for female breast tumors. (iv) ChIP-seq and inferred copy number tracks for male breast tumors. A detailed interpretation of this picture is presented in the main text. The BC hub tracks shown are based on data from (Ross-Innes et al., 2012; Severson et al., 2018).

The middle and lower parts of the image show native ChIP-seq and computationally inferred copy number tracks for selected female and male tumors. We note strongly amplified regions in female tumor 1 and male tumor 7. Overall, the two types of tracks exhibit convergent signals. However, we note a conspicuous dark black band on the left side of the viewing range, which is present in all ChIP-seq samples. Unexpectedly, this band also appears in tumors, which appear to be normal diploid in this region according to the copy number tracks, suggesting it may reflect a true cluster of ER binding peaks. This case illustrates the added value of computationally inferred copy number tracks over raw ChIP-seq tracks.

### CAGE and ChIP-seq data reflect the activity of an enhancer of the AIM2 gene

The AIM2 protein, a component of the inflammosome, reportedly suppresses cell proliferation and tumor growth in breast cancer (Chen et al., 2006). Expression of the AIM2 gene is regulated by an exapted (recycled for a new function) retroelement of the MER41 family, which contains two adjacent STAT1 binding motifs (Schmid and Bucher, 2010). These binding sites render the gene γ-interferon inducible via the JAK-STAT pathway (Chuong et al., 2016). In gastrointestinal cancers, up-regulation of AIM2 is mediated by cytokine-induced phosphorylated STAT3 (pSTAT3) (Dawson et al., 2021). ChIP-seq data for pSTA3 have recently become available for breast cancer cell lines treated with IL6 (Siersbæk et al., 2020). Since STAT 1 and STAT3 recognize similar DNA motifs, one could speculate that the aforementioned MER41 element is capable of inducing AIM2 in breast cancer via pSTAT3.

**Figure 7.**
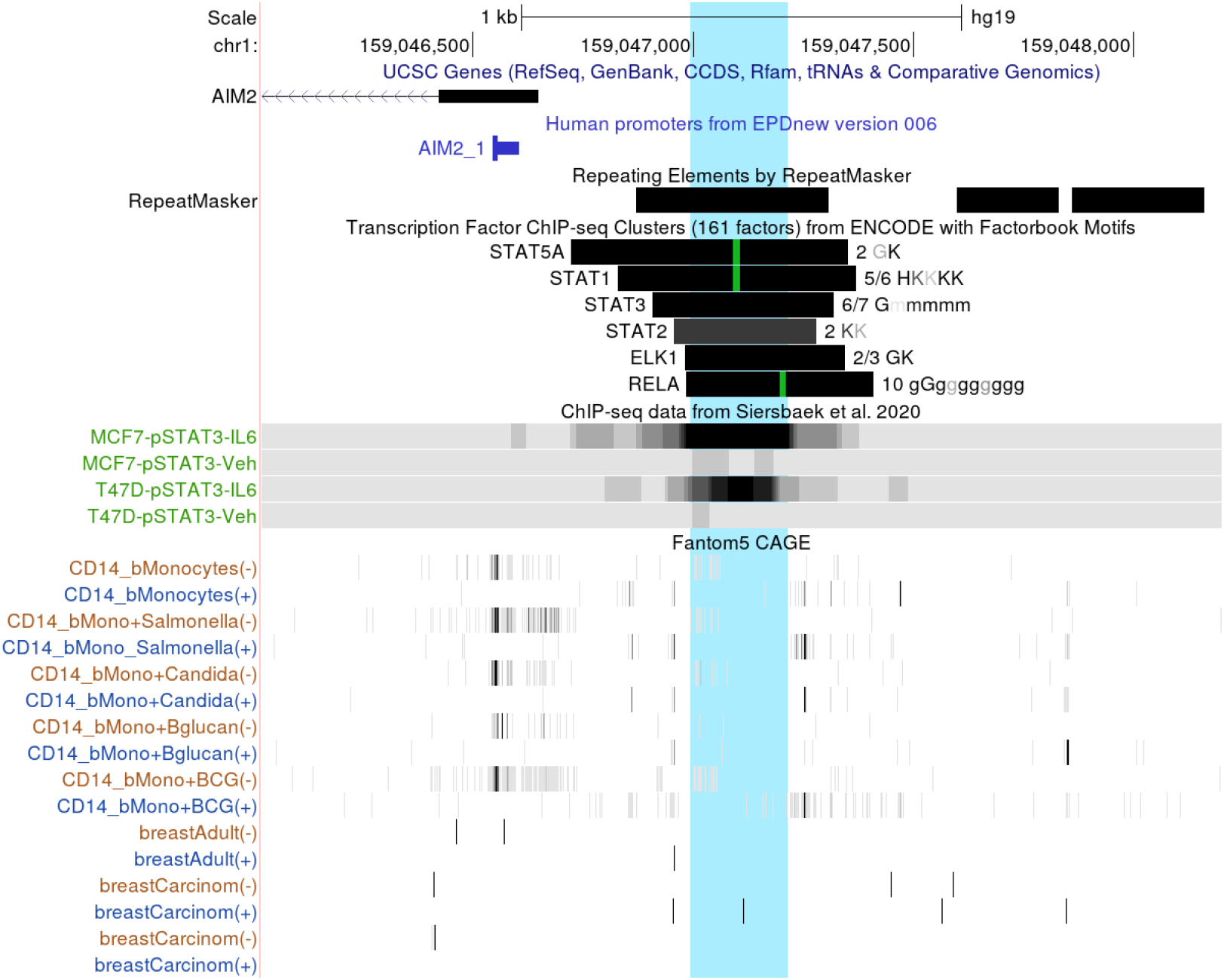
CAGE and ChIP-seq data reflect activity status of JAK-STAT inducible enhancer. The browser window is centered on an LTR MER41-derived enhancer upstream of the AIM2 gene. This screenshot is intended to illustrate the power of CAGE data in revealing the activity status of a transcriptionally active enhancer. Track sets from top to bottom: (i) Tracks from the UCSC genome browser database: UCSC genes, EPDnew promoters, merged ChIP-seq peaks from ENCODE and repeating elements from RepeatMasker; selected ENCODE peaks: all available STAT proteins, one representative member of the Ets and NFκB families: ELK1 and RELA; (ii) BC hub: ChIP-seq tracks for pSTAT3 in unstimulated and Il6 stimulated T47D cells; EPD hub: CAGE data from Fantom5 for selected tissues and cell types. A detailed interpretation of this picture is presented in the main text. The BC hub tracks shown are based on data from (Siersbæk et al., 2020). Abbreviations in track names: Veh, vehicle; BCG, Bacillus Calmette-Guérin (tuberculosis) vaccine.

The browser screenshot in Fig. 7 is centered on the MER41 element marked by a black bar in the RepeatMasker track. The ENCODE ChIP-seq cluster track indicates binding peaks for several STAT family members plus ELK1 and RELA. Highlighted in green are STAT and NFκB (RELA) DNA binding motifs. The ChIP-seq tracks from Siersbaek et al. 2020 show that this enhancer region is bound by pSTAT3 upon IL6 stimulation in two breast cancer cell lines, MCF7 and T47D.

The lower part of the image shows CAGE tracks from the EPD viewer hub, reflecting transcription initiation events. Since the MER41 element under investigation is a transcriptionally active enhancer, such data can reveal its activity status across different tissues. We note transcriptional activity in the antisense direction in CD14 blood monocytes under some conditions, in particular upon stimulation with BCG (tuberculosis) live vaccine. Conversely, in unstimulated normal and cancerous breast tissue, this enhancer appears to be silent, consistent with the pSTAT3 ChIP-seq data for breast cancer cell lines. Taken together, these observations suggest that AIM2 is silent in breast cancer cells under unstimulated conditions but induced by IL6 via pSTAT3 binding to the MER41 element.

This example was chosen in part to illustrate the value of CAGE tracks from the FANTOM5 consortium, which cover more than 500 tissues and cell types. In many situations, these are the only available data that could shed light on the activity status of an enhancer of interest.

### Work in progress and future perspectives

We make continuous efforts to expand the BC hub’s coverage, optimize the display settings and browser menu configuration, improve the sample descriptions and standardize the metadata. Some of the focus areas for improvement will be addressed in more detail below along with planned extensions and novel features.

#### Sample annotation, metadata standards and track naming

Sample annotation is a labor-intensive process, which requires domain-specific biological knowledge and currently cannot be automatized. Track description and naming is an art rather than a science as multiple conflicting objectives have to the met. All descriptive elements should be informative and understandable to as many users as possible. They should further be consistent with the nomenclature used in the corresponding journal articles and compliant with international nomenclature guidelines. Additional constraints apply to track names, which must be short in order to fit into the limited space reserved for them on the left side of the USCS genome browser window, but nevertheless unique within the track set, in order to unambiguously identify all visible tracks.

We are far from having reached the quality standards we are aiming at with regard to the objectives stated above. Several measures are taken to improve the situation. We are in the process of writing up guidelines defining annotation standards and minimal requirements for sample descriptions. This will include a list of mandatory nomenclature resources to be used. The guidelines will be posted on the web site once we have reached a satisfactory level of compliance within the BC hub. Hopefully, this will also help solicit feedback from the authors of the data. A consequence of ongoing standardization efforts is that current sample descriptions are expected to undergo frequent modifications in the near future. We apologize in advance to the reader unable to spot a track name appearing in a Figure from this paper during a live genome browser session.

#### Track signal normalization

Signal normalization, including correction for confounding effects and technical artifacts, is a pervasive and largely unsolved problem in ChIP-seq data analysis. Background information on this topic and references to published methods can be found in a recent review (Nakato and Sakata, 2021). Input controls could be useful but are not always available. Controls from pooled tumor samples can be misleading when applied to individual tumors because of tumor-specific copy number variation. These are some of the reasons why we currently present tracks as un-normalized read counts in bins of fixed length.

The track DB format offers the possibility to define a default vertical viewing range. We currently use this feature to provide customized upper limits for each track, extrapolated from the global count density over the entire genome. However, we have observed that this approach often leads to suboptimal track visualization due to local bias in read counts. Therefore, we usually overwrite the default limits via the “auto-scale to data view” option in the session links to customized views posted on the BC hub homepage. This will automatically set the viewing range to the lowest and highest track value in the current browser window. One side effect of this is that high background noise appears if there is no true signal in the selected genomic regions, see for instance the ChIP-seq tracks of the negative tumor sample in Fig. 4A. We will continue to work on this problem by evaluating novel ChIP-seq signal normalization methods as they become available.

#### Increasing data coverage

The current version of the BC hub contains only a fraction (presumably less than 50%) of all public data sets that would qualify for inclusion and be of interest to users. The bottleneck is the manual work needed for the identification of interesting data sets, quality control and sample annotation. In order to scale up this process, we envisage soliciting help from the data producers. Researchers who would like to give their own data more visibility through the BC hub are encouraged to contact us. Our data curation efforts could also benefit from feedback from the user community, including notifications of errors in the sample descriptions.

#### Extension to genome assembly hg38

We currently only support human genome assembly hg19 (GRCh37) of Feb. 2009. In the future, we plan to lift over all BC hub tracks to the newer hg38 (GRCh38) assembly. This will be a significant advance because most public tracks (including those from UCSC genome browser database) exist only for one but not the other assembly. A new hg38 version of the BC hub would thus enable users to jointly visualize tracks from the BC hub with a host of new tracks. While the “lift-over” operation to hg38 can be done automatically and swiftly, the design of new customized views for a new data environment will take some time and require creative work.

#### Support for genetic data

Currently, the BC hub offers computationally derived copy number tracks for one track set only, Severson et al. 2018 (Fig. 6). We plan to generate similar tracks for all other ChIP-seq samples in the near future. Note that some of our customized views already include third party tracks on disease associated germline variants.

What is currently missing are single nucleotide variant (SNV) tracks for tumor samples and cell lines. We have great expectations that side-by-side visualization of ChIP-seq, CNV and SNV tracks for the same tumor will help elucidate patientspecific cancer driver events at the molecular level. Providing access to cancer cell line genomes via browser viewable tracks in variant calling format (VCF) is thus high on our priority list. A rather ambitious goal would be to offer partial somatic SNV profiles extracted from patient-derived ChIP-seq and ATAC-seq samples. This seems nevertheless feasible as it works already with RNA-seq data (Coudray et al., 2018).

#### Coordination and integration with other initiatives

So far, we have developed the BC hub autonomously. However, coordination with similar efforts has now become a priority for the near future. Most urgently, we plan to register the BC hub at UCSC and EBI to enhance findability. We further plan to explore the feasibility of making our track metadata compatible with the emerging FAIRtracks standard (Gundersen et al., 2021).

## Methods

### Read mapping and import into the MGA repository

Currently all BC hub tracks were derived from short read sequencing data downloaded from GEO or ArrayExpress in FASTQ format. Reads were mapped to the human genome assembly hg19 with *Bowtie* or *Bowtie2* (Khan et al., 2018). The resulting read alignment files in SAM format were converted into SGA format using a cascade of three programs: *samtools* (Danecek et al., 2021), *bamToBed* (Quinlan, 2014) and *bed2sga* from the ChIP-Seq tools (Ambrosini et al., 2016). Procedures vary somewhat from one data set to another. The scripts used for generating each sample can be downloaded from the MGA web site to ensure reproducibility. Method details like parameters settings used for the command-line tools can be found there.

### Generation of ChIP-seq and ATAC-seq tracks

ChIP-seq and ATAC-seq tracks were generated from the corresponding SGA files in the MGA repository using *sga2wig* from the ChIP-Seq tools, and the *wigToBigWig* utility from University of California, Santa Cruz (UCSC). Replicate samples were usually merged. The “oriented” (strand-sensitive) SGA files were first “centered” using the program *chipcenter* from ChIP-seq tools. Centering means shifting the starting positions of the reads on the + and – strand by a fixed distance downstream and upstream, respectively. A shifting distance of 50, 75, 80 or 100 (depending on the experiment) was used for ChIP-seq, and 4 for ATAC-seq data. The centered SGA files were then converted into fixedStep wiggle (wig) format using step size 50 for ChIP-seq and 1 for ATAC-seq. Finally, the resulting wig files were converted to bigWig format using *wigToBigWig*.

### Generation of copy number tracks

Copy-number profiles in the Severson CNV track set were obtained using *CopywriteR* (v2.14.1) from *Bioconductor* (v3.8), a tool that detects DNA copy number alterations from off-target sequencing reads from ChIP-seq data. CopywriteR uses binned read count data as a basis for copy number detection. To correct for biases, it uses so-called “helper files” containing pre-assembled, binned (1 kb) GC-content and mapability information, which are genome assembly specific. It first removes “on-target” reads by means of a peak calling algorithm (MACS), and subsequently calculates the depth of coverage for the bins that are provided in the helper files *(CopywriteR* function).

Input to *CopywriteR* are two BAM files for each sample, a ChIP-seq sample file and a matched germline control. The BAM files were generated from FASTQ files as described in the preceding subsection. If ChIP-seq data for multiple targets were available for the same sample, those were merge. Mixed tumor input samples were used as germline controls. CNA profiles were generated for each sample using the *plotCNA* function from the output of *CopywriteR. plotCNA* generates *‘segment.Rdata’*, an R object of the DNAcopy class, which contains the segmentation values for all analyzed samples. Log2-transformed read counts were extracted from *‘segment.Rdata’* for each sample and wig files generated via *ad hoc* R and Perl scripts. The *wigToBigWig* utility from UCSC was then used to generate bigWig track files.

### QC plots

Strand correlation plots were generated as described previously (Ambrosini et al., 2016). The motif enrichment plots were produced with programs from ChIP-seq tools and precomputed match lists for TF binding motifs available from the MGA repository. These match lists were generated with the PWMScan workflow (Ambrosini et al., 2018), using the following position frequency matrices from JASPAR (Castro-Mondragon et al., 2022): MA0112.3 for ESR1 (ER), MA0007.3 for AR, MA0113.3 for NR3C1 (GR), MA0148.4 for FOXA1, MA0144.2 for STAT3, MA0102.4 for CEBPA, MA0139.1 for RAD21/CTCF and MA0103.3 for ZEB1, and from HOCOMOCO (Kulakovskiy et al., 2018): PRGR_HUMAN.H11MO.0.A for PGR (PR) and GRHL2_HUMAN.H11MO.0.A for GRHL2.

The motif matches falling into annotated repeat regions of the UCSC RepeatMasker track (shown in Fig. 7) were first filtered out with the program *counts_filter*. A centered version of the SGA file containing read map position was then generated with *chipcenter* using a shifting distance of 75 bp. The resulting file was used to determine the read coverage around each motif match with *chipscore*. The top 1000 motif matches in terms of read coverage were selected for further analysis. Strand-separated aggregation plots of ChIP-seq reads around selected motifs were generated with *chipcor* and combined in a single plot using R software. Details of the procedures including the parameters uses for each program can be found in the corresponding script directories (hyperlinked to the MGA series documentation pages).

